# A mechanistic model of linkage analysis in allohexaploids

**DOI:** 10.1101/035139

**Authors:** Huan Li, Xuli Zhu, Qin Yan, Ke Mao, Rongling Wu

## Abstract

Despite their pivotal role in agriculture and biological research, polyploids, a group of organisms with more than two sets of chromosomes, are very difficult to study. Increasing studies have used high-density genetic linkage maps to investigate the genome structure and function of polyploids and to identify genes underlying polyploid traits. However, although models for linkage analysis have been well established for diploids, with some essential modifications for tetraploids, no models have been available thus far for polyploids at higher ploidy levels. The linkage analysis of polyploids typically requires knowledge about their meiotic mechanisms, depending on the origin of polyplody. Here we describe a computational modeling framework for linkage analysis in allohexaploids by integrating their preferential chromosomal-pairing meiotic feature into a mixture model setting. The framework, implemented with the EM algorithm, allows the simultaneous estimates of preferential pairing factors and the recombination fraction. We investigated statistical properties of the framework through extensive computer simulation and validated its usefulness and utility by analyzing a real data from a full-sib family of allohexaploid persimmon. Our attempt in linkage analysis of allohexaploids by incorporating their meiotic mechanism lays a foundation for allohexaploid genetic mapping and also provides a new horizon to explore allohexaploid parental kinship.

## INTRODUCTION

Polyploidy is an important force for the evolution of plants (Otto and Whitton 2000; Soltis and Soltis 2000). It was estimated that 70 – 80% of angiosperms are polyploids or experienced phases of polyploids during their evolutionary process (Lewis 1979; Masterson 1994). Many crops, such as wheat, sugarcanes, potato, cotton and canola, are polyploids, which play a central role in agriculture (Leitch and Leitch 2008). Polyploids can be classified into two types, i.e., allopolyploids, whose chromosomes are composed of distinct genomes through interspecific hybridization, and autopolyploids, in which the chromosome doubling of genetically similar genomes is due to the fusion of unreduced gametes (Müntzing 1936; Soltis and Soltis 2000; Soltis et al. 2004; Gaeta and Pires 2010). Of all ployploids, more than 75% are found to be allopolyploids (Soltis and Soltis 2009). A growing body of evidence indicates that polyploids have great advantages in response to selection and adaption partly through increased rates of meiotic recombination (Soltis and Soltis 1999; Grant 2004; Comai 2005; Chen 2010; Pecinka et al. 2011).

The nature of polyploids can be depicted through how chromosomes pair at meiosis. According to this criterion, polyploids can be sorted into bivalent polyploids, multivalent polyploids and mixed polyploids (Comai 2005). In general, extreme allopolyploids present bivalent formation in which more similar chromosomes are expected to have higher pairing frequencies than less similar chromosomes, a phenomenon which can be described by the preferential pairing factor (Sybenga 1988). On the other hand, extreme autopolyploids are pervaded by multivalent formation in which more than two chromosomes pair at a time, resulting in the appearance of two sister chromatids into the same gamete, called double reduction (Hauber et al. 1999). Mixed polyploids with both bivalent and multivalent formation are confounded by both preferential pairing and double reduction (Wu et al. 2004; Burke et al. 2015).

The past two decades have witnessed increasing studies of linkage mapping in polyploids (Ripol et al. 1999; Luo et al. 2001a,b; Kriegner et al. 2003; McCord al. 2011; Hackett et al. 2013, 2014; Monden et al. 2015; Bourke et al. 2015). Linkage maps are constructed on the basis of segregation and transmission of genes into the next progeny generation. Due to some unique cytological phenomena during meiosis, e.g., double reduction and preferential pairing, statistical models for linkage analysis in polyploids should be qualitatively more complex than those in diploids. This complexity has led to tremendous development of powerful statistical models for linkage analysis and QTL mapping in tetraploid (Hackett et al. 1998, 2013; Luo et al. 2001; Rehmsmeier 2013). Sybenga (1965, 1966) recognized the event of unequal paring probability during chromosome synapsis and developed mathematical models to describe different chromatid paring probabilities in polyploids. By taking bivalent and multivalent pairing formations into account during meiosis, Wu and group developed a series of models for linkage analysis, map construction and QTL mapping (Wu et al. 2001a,b, 2002; Ma et al. 2002; Lu et al. 2013; Yang et al. 2013; Xu et al. 2014a,b). The phenomenon of double reduction was also considered in Luo et al.’s (2001a,b) autotetraploid model and Rehmsmeier’s (2013) computational model. More recently, Li et al. (2010) developed a specialized EM algorithm for QTL mapping in multivalent tetraploids, which may impact in the field of polyploid QTL mapping. Overcoming the drawback of two-point linkage analysis, Yang et al. (2013) and Lu et al. (2013) developed a three-point linkage analysis model which can not only accurately estimate the linkage between loci, but also detect genetic interference throughout the genome.

While all these polyploidy linkage models are focused on tetraploids, there is still a gap in the model development of linkage analysis in hexaploids despite their significant importance in agriculture and biology (Monden et al. 2015). In this article, we describe and assess a statistical model that embeds preferential chromosomal-pairing within the framework of linkage analysis in allohexaploids. A considerable body of evidence shows that chromosome pairing occurs between homeologues during meiosis and homeologous recombination plays an important role in chromosomal rearrangements (Nicolas et al. 2007; Lim et al. 2008; Gaeta and Pires 2010; Xiong et al. 2011). The new model allows us to simultaneously estimate the preferential pairing factor and recombination fraction between any pair of molecular markers. We outline a detailed procedure to test the significance of these two parameters, facilitating the studies of allohexaploid genome structure and organization. The model offers a useful tool for linkage mapping and population genetic studies in allohexaploids.

## The Model

### Preferential pairing factor

The probability, with which more similar chromosomes pair more frequently than less similar chromosomes, is defined as the preferential pairing factor (Sybenga 1966). For an allopolyploid system exhibiting bivalent formation, it is expected that the preferential pairing factor influences gamete formation and frequencies. Consider a heterozygous allohexaploid derived from the chromosomal combination of three distinct diploid genomes **A**, **B** and **C**.

Six sets of chromosomes in this allohexaploid are labeled as **1**, **2**, **3**, **4**, **5** and **6**, respectively. Assume that chromosomes **1** and **2** are homologous, as are chromosomes **3** and **4**, as well as chromosomes **5** and **6**. Under bivalent pairing, there are a total of 15 possible pair-wise combinations among six chromosomes, expressed as **1** and **2**, **1** and **3**,…, **5** and **6**. Denote the preferential pairing factor as *θ*_1_ between chromosome **1** and **2**, *θ*_2_ between chromosome **3** and **4**, and *θ*_3_ between chromosome **5** and **6**. Thus, the frequencies of pairing of any two chromosomes are derived as

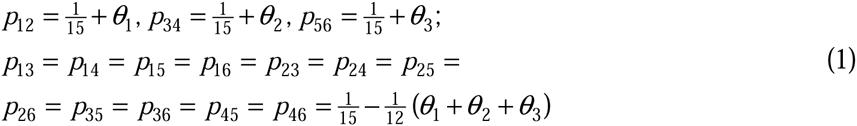

where the subscripts stand for 15 possible chromosome pairs.

For an allohexaploid genotype **123456**, its bivalent pairing takes place in three different ways: fully preferential pairing, partially preferential pairing and no preferential pairing, each of which is derived from a particular chromosomal-pairing configuration that occurs with a different frequency and generates different groups of triploid gametes at meiosis (Table 1). A chromosomal-pairing configuration is defined by using || to separate pairing chromosomes. For example, **12||34||56** is a chromosomal-pairing configuration in which chromosome 1 pairs with 2, 3 with 4 and 5 with 6. Expressions for the frequencies of chromosomal-pairing configurations are given in supplementary materail 1. Each chromosomal-pairing configuration produces eight triploid gametes, leading to 15 × 8 = 120 gametes in total. Virtually, these gametes are distinguished by 20 types, i.e., 123, 124,…, 456, whose frequencies are expressed as *g*_123_, *g*_124_,…, *g*_456_, respectively.

**Table 1.** Types and frequencies of chromosomal-pairing configurations, as well as triploid gametes each configuration produces during meiosis, for an allohexaploid with six chromosomes labeled as **1, 2, 3, 4, 5**, and **6**.

| No. | Pairing | Configuration | Freq. | Gamete |
| --- | --- | --- | --- | --- |
| 1 | Fully preferential | **12 \|\| 34 \|\| 56** | $f_1$ | **135 136 145 146 235 236 245 246** |
| 2 | Partially preferential | **12 \|\| 35 \|\| 46** | $f_2$ | **134 136 145 156 234 236 245 256** |
| 3 | Partially preferential | **12 \|\| 36 \|\| 45** | $f_3$ | **134 135 146 156 234 235 246 256** |
| 4 | Partially preferential | **15 \|\| 34 \|\| 26** | $f_4$ | **123 124 136 146 235 245 356 456** |
| 5 | Partially preferential | **16 \|\| 34 \|\| 25** | $f_5$ | **123 124 135 145 236 246 356 456** |
| 6 | Partially preferential | **13 \|\| 24 \|\| 56** | $f_6$ | **125 126 145 146 235 236 345 346** |
| 7 | Partially preferential | **14 \|\| 23 \|\| 56** | $f_7$ | **125 126 135 136 245 246 345 346** |
| 8 | No preferential | **13 \|\| 25 \|\| 46** | $f_8$ | **124 126 145 156 234 236 345 356** |
| 9 | No preferential | **13 \|\| 26 \|\| 45** | $f_9$ | **124 125 146 156 234 235 346 356** |
| 10 | No preferential | **14 \|\| 25 \|\| 36** | $f_{10}$ | **123 126 135 156 234 245 345 456** |
| 11 | No preferential | **14 \|\| 26 \|\| 35** | $f_{11}$ | **123 125 136 156 234 245 346 456** |
| 12 | No preferential | **15 \|\| 23 \|\| 46** | $f_{12}$ | **124 126 134 136 245 256 345 356** |
| 13 | No preferential | **15 \|\| 24 \|\| 36** | $f_{13}$ | **124 126 134 136 245 256 345 356** |
| 14 | No preferential | **16 \|\| 23 \|\| 45** | $f_{14}$ | **124 125 134 135 246 256 346 356** |
| 15 | No preferential | **16 \|\| 24 \|\| 35** | $f_{15}$ | **123 125 134 145 236 256 346 456** |

### Meiotic chromosomal segregation

Our model focuses on an allohexaploid that undergoes only bivalent pairing during meiosis. Suppose that a heterozygous allohexaploid line is crossed with a homozygous line to generate a pseudo-test backcross in which the genotypes of the progeny are consistent with the genotypes of gametes produced by the heterozygous parent. Assume that there are two fully informative markers, **A** and **B**, which are both heterozygous in one parent but homozygous in another parent. We denote six different alleles as *a*_1_,…, *a*_6_ at marker **A** and *b*_1_,…,*b_6_* at marker **B**. Nonalleles at the two markers are located in six chromosomes with 6 × 5 × 4 × 3 × 2 × 1 = 720 possible linkage phases, expressed as

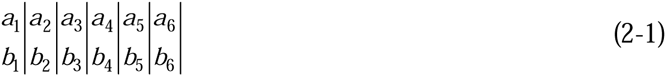

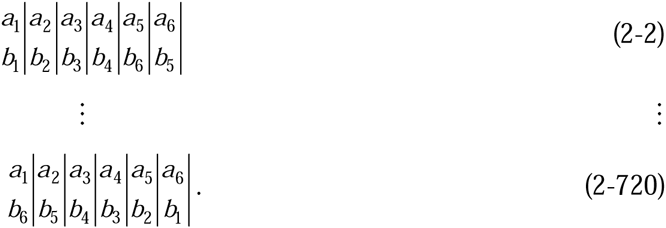

A heterozygous allohexaploid with one particular linkage phase above has 15 possible chromosomal-pairing configurations. For linkage phase (2-1), these configurations can be expressed, in the order shown in Table 1, as

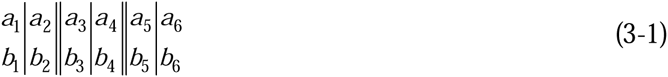

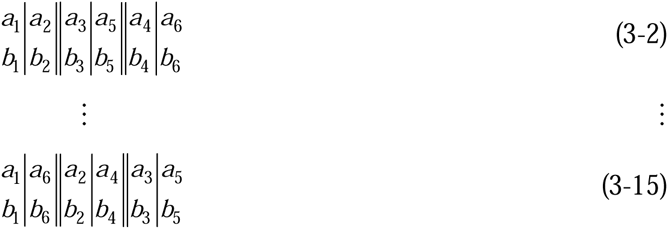

where we assume that chromosomes **1** and **2**, **3** and **4**, and **5** and **6** are each homologous. Under fully preferentially pairing (3-1), marker **A** produces eight triploid gametes *a*_1_*a*_3_*a*_5_, *a*_1_*a*_3_*a*_6_, *a*_1_*a*_4_*a*_5_, *a*_1_*a*_4_*a*_6_, *a*_2_*a*_3_*a*_5_, *a*_2_*a*_3_*a*_6_, *a*_2_*a*_4_*a*_5_, and *a*_2_*a*_4_*a*_6_ with the same thing as marker **B**. Let *r* denote the recombination fraction between markers **A** and **B**. According to the number of crossover between markers **A** and **B**, meiotic gametes fall into 4 categories: no crossover, one crossover, two crossovers and three crossovers, with frequencies denoted as *p*_0_*, p*_2_*, p*_2_*, p*_3_, and *p*_4_, respectively, which are expressed, in terms of *r*, as follows:

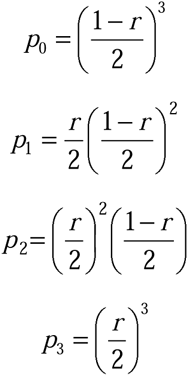

By combining three haploid gametes each from a different chromosome, triploid gametes are generated. Table 1 lists all possible triploid gametes and their probabilities produced under fully preferentially pairing (3-1) of linkage phase (2-1).

### Likelihood, estimation and tests

Consider a full-sib family derived from two allohexaploid parents in which markers may be segregating in two manners. One is the intercross segregation at which both parents are heterozygous. The second is the testcross segregation at which one parent is heterozygous whereas the second is homozygous. As the demonstration of model derivation, we consider two testcross markers by crossing the parents

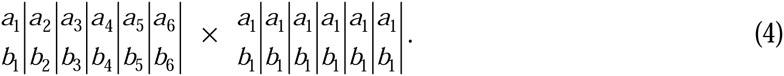

Let 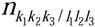 denote the observation of progeny with triploid gamete genotype *k*_1_*k*_2_*k*_3_ (*k*_1_ *< k*_2_ < *k*_3_ = 1,…, 6) at marker **A** and triploid gamete genotype *l*_1_*l*_2_*l*_3_ (*l*_1_ *< l*_2_ < *l*_3_ = 1,…, 6) at marker **B** derived from the heterozygous parent. Correspondingly, the probability of a two-marker triploid gamete genotype is denoted as 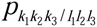. Then, we formulate a likelihood for observed genotype data, expressed as

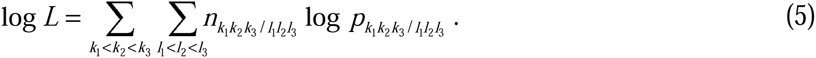

If the heterozygous parent has a certain chromosomal-pairing configuration (Table 1), 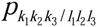 only contains the unknown recombination fraction *r*. By maximizing the likelihood (5), the maximum likelihood estimate (MLE) of *r* can be obtained by an explicit expression.

In practice, the chromosomal-pairing configuration of an allohexaploid is unknown. However, given that it presents a mix of 15 possible configurations (Table 1), for the same triploid gamete genotype, we can derive 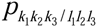 as a sum of its frequencies (determined by *p*_0_*, p*_1_*, p*_2_, or *p*_3_) weighted by chromosomal-pairing configuration frequencies *f*_1_,…,*f*_15_. With the derived 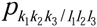, the likelihood (5) was reformulated as a mixture model. The EM algorithm (given in supplementary material 2) can be implemented to obtain the estimates of preferential pairing factors *θ*_1_*, θ*_2_ and *θ*_3_ and recombination fraction *r*. It is shown that the estimation of *r* can be obtained from a closed form.

For a practical marker data, we do not know the linkage phase and chromosomal homology of an allohexaploid a priori. We can infer these two uncertainties from the marker genotype data to obtain the correct estimate of recombination fraction *r*. First, an allohexaploid may have 180 types of linkage phases over two markers each with six different alleles. Second, there are 15 possible homologuous relationships among six chromosomes, i.e.,

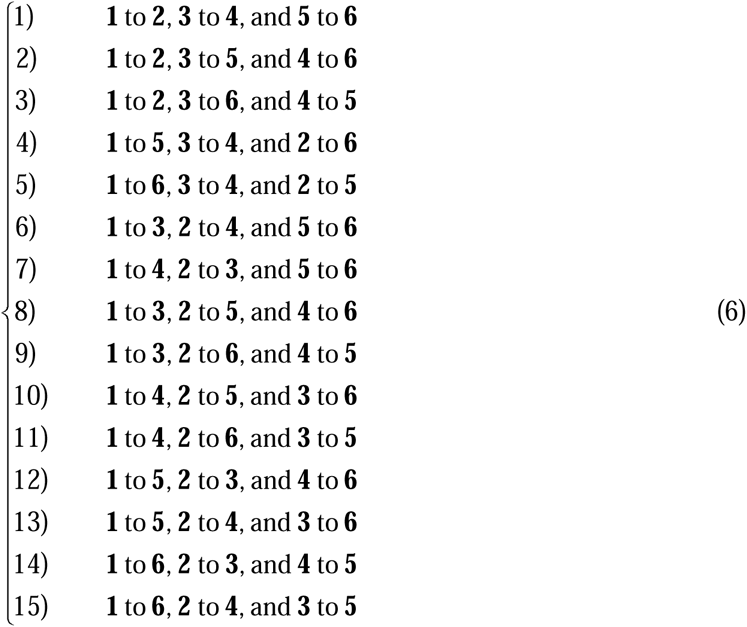

Thus, we need to choose an optimal combination of linkage phase and chromosomal homology from a total of 180 × 15 = 2700 possibilities. By obtaining the corresponding 2700 plug-in likelihood values, we select the largest one that corresponds to the optimal combination fromn which the MLEs of preferential pairing factors and recombination fraction can be obtained.

After the unknown parameters (*θ*_1_, *θ*_2_ and *θ*_3_ and *r*) are estimated under an optimal linkage phase and homology, we formulate a series of hypotheses to test if each of these parameters is significant. These tests include those of whether the recombination fraction is different from 0.5, whether there is no preferential pairing between more similar chromosomes during meiosis, i.e. *θ*_1_ = *θ*_2_ = *θ*_3_ = 0, and whether one of the preferential pairing does not exist between a particular pair of more similar chromosomes during meiosis, i.e. *θ*_1_ *=* 0 or *θ*_2_ *=* 0 or *θ*_3_ *=* 0. To test these hypotheses, we need to calculate the log-likelihood ratios from the likelihoods under the null and alternative hypotheses and compare it against a critical value obtained from a chi-square distribution with three or one degree of freedom.

### Linkage model for partially informative markers

In the preceding section, a procedure was described for linkage analysis of fully informative markers (with six distinct alleles at each marker) in allohexaploids, but a consideration should be taken for those partially informative markers which have multiple copies of the same alleles at one or two markers. For partially informative markers, a mixture likelihood constructed under a particular allelic configuration can be similarly constructed, but with a more complex structure due to their inconsistency between observed genotypes and real configurations. For example, a five-allele genotype observed as *a*_1_*a*_2_*a*_3_*a*_4_*a*_5_, may have five possible configurations; i.e., *a*_1_|*a*_1_|*a*_2_|*a*_3_|*a*_4_|*a*_5_|, *a*_1_|*a*_2_|*a*_2_|*a*_3_|*a*_4_|*a*_5_|, *a*_1_|*a*_2_|*a*_3_|*a*_3_|*a*_4_|*a*_5_|, *a*_1_|*a*_2_|*a*_3_|*a*_4_|*a*_4_|*a*_5_|, and *a*_1_|*a*_2_|*a*_3_|*a*_4_|*a*_5_|*a*_5_|, but for a four-allele genotype observed as *b*_1_*b*_2_*b*_3_*b*_4_, it has as many as 10 possible configurations, such as *b*_1_|*b*_1_|*b*_1_|*b*_2_|*b*_3_|*b*_4_|, *b*_1_|*b*_1_|*b*_2_|*b*_2_|*b*_3_|*b*_4_|, *b*_1_|*b*_1_|*b*_2_|*b*_3_|*b*_3_|*b*_4_|, *b*_1_|*b*_1_|*b*_2_|*b*_3_|*b*_4_|*b*_4_|, *b*_1_|*b*_2_|*b*_2_|*b*_2_|*b*_3_|*b*_4_|, *b*_1_|*b*_2_|*b*_2_|*b*_3_|*b*_3_|*b*_4_|, *b*_1_|*b*_2_|*b*_2_|*b*_3_|*b*_4_|*b*_4_|, *b*_1_|*b*_2_|*b*_3_|*b*_3_|*b*_3_|*b*_4_|, *b*_1_|*b*_2_|*b*_3_|*b*_3_|*b*_4_|*b*_4_|, and *b*_1_|*b*_2_|*b*_3_|*b*_4_|*b*_4_|*b*_4_|. A three- or two-allele genotype has 10 and five different configurations, respectively. An extra difficulty for linkage analysis of partially informative markers lies in the estimation of the probability at which each allelic configuration occurs and then the determination of the most likely configuration.

For two fully informative markers, there are 15 distinguishable chromosomal homologies (6). But some of these homologies are collapsed into the same group for partially informative markers. Also, in such a case, the triplotypes of gametes are collapsed because of indistinguishable types of recombinants and non-recombinants. These two types of collapses together make it more difficult to estimate the preferential pairing factors and recombination fraction the EM algorithm. We have derived a general procedure for estimating these two parameters and testing their significance when two markers are partially informative. It should be noted that, for fully informative markers and five- and four-allele partially informative markers, all three preferential pairing factors (*θ*_1_, *θ*_2_ and *θ*_3_) that determine chromosomal pairing types can be estimated, but because of reduced degrees of freedom, only one and two preferential pairing factors can be estimated for two- and three-allele partially informative markers, respectively.

## Results

### Computer simulation

We performed Monte Carlo simulation to investigate the statistical properties of the allohexaploid linkage analysis model. The simulation experiments were designed to reflect ranges of the preferential pairing factor *θ*_1_ *=* 0.15, 0.10 and 0.05 and recombination fraction *r* = 0.05, 0.15 and 0.30. The data of marker genotypes were simulated for two testcross markers in a full-family of size *N* = 100, 200, or 400 by assuming a particular linkage phase for the two markers and chromosomal homology.

Table 3 shows the results about parameter estimation from 1000 simulation replicates by the new model. It appears that a small sample size 100 can provide reasonably good estimates of the preferential pairing factor and recombination fraction, but the accuracy and precision of parameter estimates increase dramatically with increasing sample size. The linkage of two highly linked markers (*r* = 0.05) can be better estimated than that of two loosely linked markers (*r* = 0.30). The preferential pairing factors can be well estimated, not depending on the degree of linkage between two markers. The power to detect the correct linkage phase and homology is quite high. This is not surprising because the segregation of marker genotypes is very sensitive to the pattern of linkage phase and homology.

We performed an additional simulation study to examine the power of detecting the linkage and preferential pairing factors. The empirical power of the detection of these parameters was calculated by considering different sample sizes and different degrees of linkage (Table 2). In general, the power to detect the linkage is very high, which is not surprised because the MLE of the recombination fraction was based on an explicit expression. The power to jointly detect all possible preferential pairing is also very high, but reduced for the detection of individual preferential pairing. When the preferential pairing factor is low (say 0.05), the power to detect it becomes very low especially when sample size is modest (100). As shown in Table 2, if the preferential pairing factor is about 0.10, sample size of 200 – 400 is required to detect its occurrence. When the preferential pairing factor is low, e.g., 0.05, sample size of over 400 should be used.

**Table 2.** The probabilities of all possible triploid gametes for markers **A** and **B** produced by an phase-known allohexaploid, diagrammed as 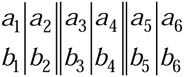, under fully preferentially pairing.

| Marker A |  |  |  |  |  |  |  |  |
| --- | --- | --- | --- | --- | --- | --- | --- | --- |
| Marker B | $a_1\vert a_3\vert a_5\vert $ | $a_1\vert a_3\vert a_6\vert $ | $a_1\vert a_4\vert a_5\vert $ | $a_1\vert a_4\vert a_6\vert $ | $a_2\vert a_3\vert a_5\vert $ | $a_2\vert a_3\vert a_6\vert $ | $a_2\vert a_4\vert a_5\vert $ | $a_2\vert a_4\vert a_6\vert $ |
| $b_1\vert b_3\vert b_5\vert $ | $p_0$ | $p_1$ | $p_1$ | $p_2$ | $p_1$ | $p_2$ | $p_2$ | $p_3$ |
| $b_1\vert b_3\vert b_6\vert $ | $p_1$ | $p_0$ | $p_2$ | $p_1$ | $p_2$ | $p_1$ | $p_3$ | $p_2$ |
| $b_1\vert b_4\vert b_5\vert $ | $p_1$ | $p_2$ | $p_0$ | $p_1$ | $p_2$ | $p_3$ | $p_1$ | $p_2$ |
| $b_1\vert b_4\vert b_6\vert $ | $p_2$ | $p_1$ | $p_1$ | $p_0$ | $p_3$ | $p_2$ | $p_2$ | $p_1$ |
| $b_2\vert b_3\vert b_5\vert $ | $p_1$ | $p_2$ | $p_2$ | $p_3$ | $p_0$ | $p_1$ | $p_1$ | $p_2$ |
| $b_2\vert b_3\vert b_6\vert $ | $p_2$ | $p_1$ | $p_3$ | $p_2$ | $p_1$ | $p_0$ | $p_2$ | $p_1$ |
| $b_2\vert b_4\vert b_5\vert $ | $p_2$ | $p_3$ | $p_1$ | $p_2$ | $p_1$ | $p_2$ | $p_0$ | $p_1$ |
| $b_2\vert b_4\vert b_6\vert $ | $p_3$ | $p_2$ | $p_2$ | $p_1$ | $p_2$ | $p_1$ | $p_1$ | $p_0$ |

To investigate how the model performs for the linakge analysis of partially informative markers, a simulation study was carried out by hypothosizing two markers **A:** *a*_1_*a*_2_*a*_3_*a*_3_*a*_4_*a*_5_ and **B:** *b*_1_*b*_2_*b*_2_*b*_3_*b*_4_*b*_4_ of different strengths of linkage (*r* = 0.05, 0.15 and 0.30). The preferential pairing factors were assumed as *θ*_1_=0.15, *θ*_2_=0.10 and *θ*_3_=0.05. We assume a backcross design of different sample sizes n = 100, 200 and 400. Table 5 gives the estimates of the preferential pairing factors and recombination fraction under these scenarios. The model can provide reasonably accurate estimates of these parameters. As expected, the precision of parameter estimation increases with increasing sample size. The estimation of the preferential pairing factors is generally independent of the degree of linkage. For partially informative markers, the power to correctly detect both allelic configuration and chromosomal homology is about 0.5 with a modest sample size. However, we noted that power would increase to > 0.95 if the selected allelic configuration differs by one chromosome from the true configuration. Table 6 lists the power of linkage detection and preferential pairing detection under different simulation scenarios. In general, all the power is quite high even for a modest sample size (100), but to detect preferential pairing, a large sample size (300) is needed if the markers are loosely linked.

**Table 3.** MLEs of preferential pairing factors and recombination fraction and their standard deviations over 1000 simulation replicates estimated from a simulated marker data generated by a phase- and homology-known allohexaploid under different simulation scenarios by changing the recombination fraction and sample size. The power of detecting correct linkage phase and correct homology at the same time is also given.

| Simulation Scenario | $\theta_1$<br>(0.15) | $\theta_2$<br>(0.10) | $\theta_3$<br>(0.05) | $r$ | Power |
| --- | --- | --- | --- | --- | --- |
| $r = 0.05$ | | | | | |
| 100 | 0.146±0.093 | 0.103±0.093 | 0.061±0.080 | 0.050±0.012 | 1 |
| 200 | 0.148±0.069 | 0.100±0.065 | 0.058±0.061 | 0.051±0.009 | 1 |
| 400 | 0.150±0.051 | 0.103±0.047 | 0.052±0.044 | 0.050±0.006 | 1 |
| $r = 0.15$ | | | | | |
| 100 | 0.146±0.093 | 0.103±0.093 | 0.061±0.080 | 0.151±0.020 | 1 |
| 200 | 0.148±0.069 | 0.100±0.065 | 0.058±0.061 | 0.150±0.014 | 1 |
| 400 | 0.150±0.051 | 0.103±0.047 | 0.052±0.044 | 0.150±0.010 | 1 |
| $r = 0.30$ | | | | | |
| 100 | 0.146±0.093 | 0.103±0.093 | 0.061±0.080 | 0.301±0.029 | 0.997 |
| 200 | 0.148±0.069 | 0.100±0.065 | 0.058±0.061 | 0.300±0.018 | 1 |
| 400 | 0.150±0.051 | 0.103±0.047 | 0.052±0.044 | 0.301±0.014 | 1 |

**Table 4.**
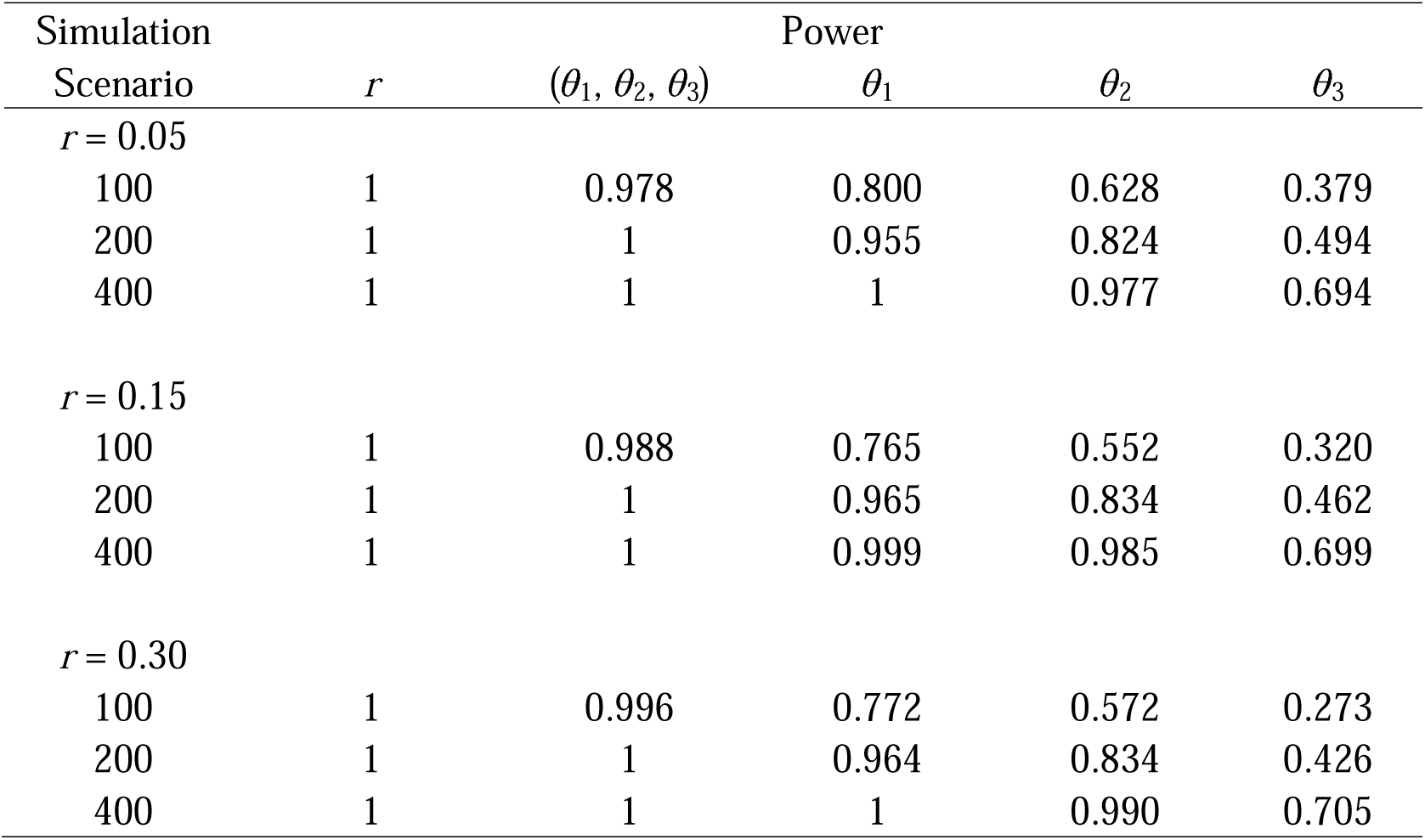
Empirical power to detect the linakge and three preferential pairing factors under different simulation scenarios by changing the recombination fraction and sample size.

**Table 5.**
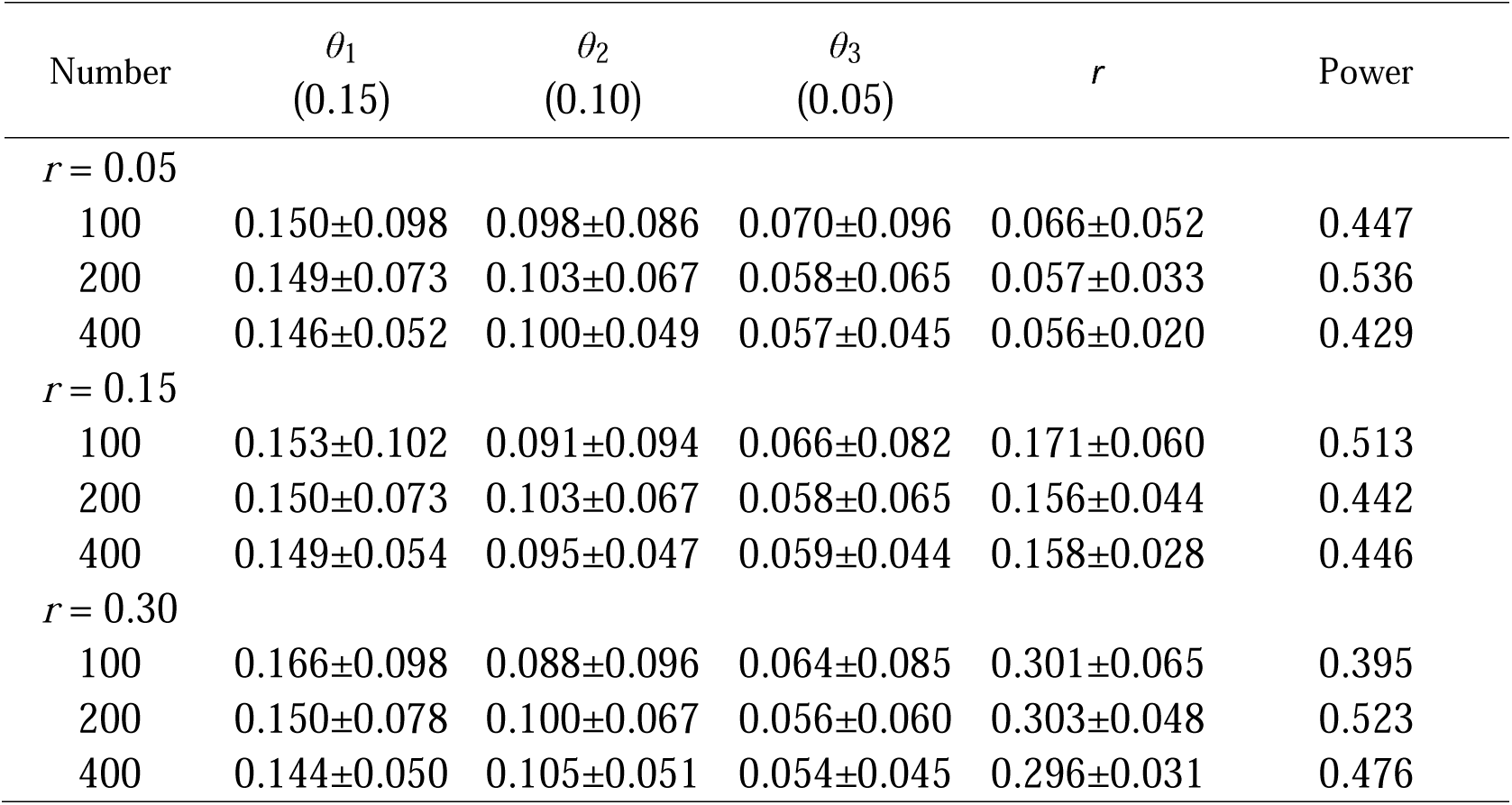
MLE of preferential pairing factors and recombination fraction and their standard deviation over 1000 simulation replicates from a simulated marker data. The power of detecting correct linkage phase and correct homology at the same time is also given.

**Table 6.** Power to detect whether the estimated parameters are significant under different hypotheses.

| Number | Power |  |  |  |  |
| --- | --- | --- | --- | --- | --- |
| | $r=0.5$ | $\theta_1=\theta_2=\theta_3=0$ | $\theta_1=0$ | $\theta_2=0$ | $\theta_3=0$ |
| $r = 0.05$ | | | | | |
| 100 | 1 | 0.992 | 0.957 | 0.886 | 0.742 |
| 200 | 1 | 1 | 0.989 | 0.972 | 0.745 |
| 400 | 1 | 1 | 1 | 0.997 | 0.926 |
| $r = 0.15$ | | | | | |
| 100 | 1 | 0.981 | 0.902 | 0.876 | 0.723 |
| 200 | 1 | 1 | 0.993 | 0.986 | 0.842 |
| 400 | 1 | 1 | 1 | 0.992 | 0.913 |
| $r = 0.30$ | | | | | |
| 100 | 0.925 | 0.981 | 0.952 | 0.883 | 0.779 |
| 200 | 1 | 1 | 0.983 | 0.975 | 0.732 |
| 400 | 1 | 1 | 1 | 0.983 | 0.862 |

### Worked example

Currently, we have a small real dataset to test the usefulness of our model. A full-sib family of persimmon was derived the cross between an allohexaploid tree (male) and a diploid tree (female) at Shandong Research Institute of Pomology, Taishang, China. The family contains 106 progeny, genotyped for several dozens of SSR markers screened from published EST-SSR primers. We analyzed four randomly chosen markers, whose mating types are detected, on the basis of Mendelian segregation law, as

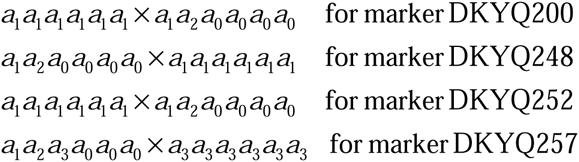

where some alleles cannot be determined precisely as *a*_1_ or *a*_2_, which are denoted by *a*_0_. Our model was equipped with a function to discern these alleles for the accurate estimation of the linkage and preferential pairing factor.

We performed pair-wise linkage analysis by first determining the most likely linkage phase and homology under which the recombination fraction and preferential pairing factors were estimated, with results given in Table 7. By a re-sampling approach, the standard errors of each estimate were obtained. The estimated recombination fractions between these two markers range from 0.056 to 0.134, showing mutual highly linked relationships. Because a few number of alleles at each marker, we can only estimate a couple of preferential pairing factors. It is interesting to see that preferential pairing does occur among chromosomes in the allohexaploid persimmon, which suggests that this hexaploid woody plant probably has experienced the combination of distinct genomes through interspecific hybridization. Different values of the preferential pairing factors estimated from different marker pairs may indicate varying degrees of relatedness among different regions of chromosomes.

**Table 7.**
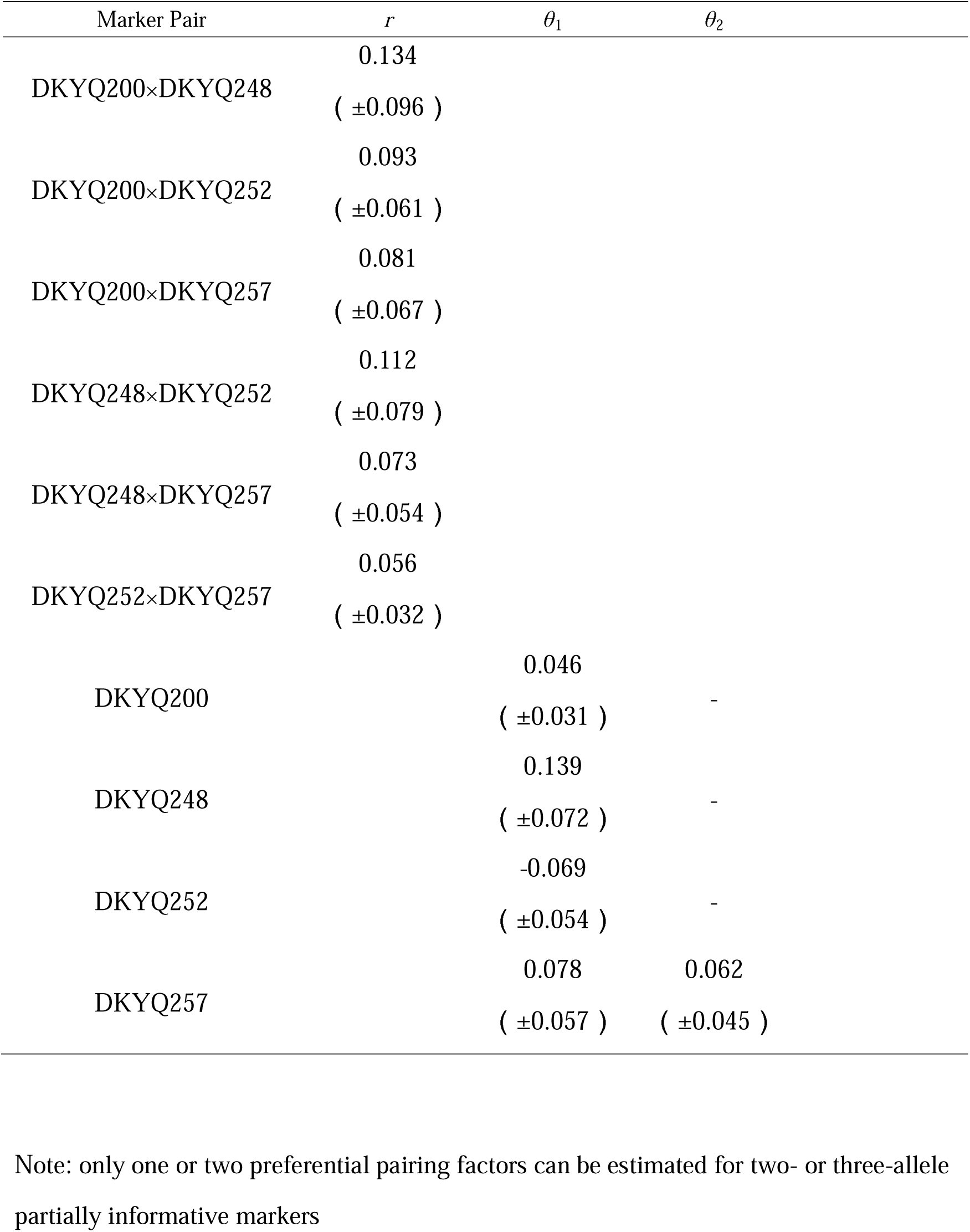
Estimates of the recombination fraction and preferential pairing factors among four SSR markers genotyped for a full-sib family of persimmon.

## Discussion

Current statistical models are mainly focused on linkage analysis for diploids (Lander and Green 1987; Stam 1993; Maliepaard et al. 1997; Wu et al. 2002). Many models for linkage analysis of polyploids are generally borrowed from diploids, which may produce misleading results because polyploids undergo qualitatively different meiotic mechanisms from diploids. For example, the frequencies of gamete formation are not only influenced by the recombination fraction, but also influenced by the relative frequencies of different chromosome pairing mechanisms, such as preferential pairing that has a widespread occurrence in allopolyploids (Sybenga 1965, 1966, 1988) and double reduction in autopolyploids (Luo et al. 2004; Wu and Ma 2005). The preferential pairing factor is an important parameter that describes the cytological characteristic of allopolyploids thought to play a key role in plant evolution. Sybenga (1992, 1998) used the preferential pairing factor to describe the homology in allopolyploids. Wu et al. (2001a) proposed that the preferential pairing factor could explain the difference between pairing formation derived from bivalent and multivalent pairings. Here, we describe and assess a model for allohexaploid linkage analysis incorporating the preferential pairing factor. Our model built upon preferential pairing and chromosomal homology shows good power to obtain more realistic results than existing linkage analysis models.

The statistical model proposed can simultaneously estimate the recombination fraction and preferential pairing factor in allohexaploids. Our model can handle the linkage of any types of markers, such as testcross markers (at which only one of the two parents is heterozygous) and intercross markers (at which both parents are heterozygous), segregating in a full-sib family of two heterozygous parents. Simulation studies were performed to investigate the statistical behavior of our model. It was found that the model displays high precision for estimating the recombination fraction and preferential pairing factor over a range of sample sizes and parameter values. By estimating the preferential pairing factor, the model helps geneticist to determine whether a particular allohexaploid undergoes preferential pairing or random pairing during meiosis and relate this information to understand the evolutionary diversity of polyploids (Nicolas et al. 2007; Lim et al. 2008; Gaeta and Pires 2010; Xiong et al. 2011).

Existing linkage analysis models for polyploids are mainly focused on triploids and tetraploids, with hexaploids being never touched before. Our model derived here fills a gap in this area. We provided a general framework for allohexaploid linkage analysis and its principle can be extended to octoploid and dexaploid species, but their increasing complexity of model derivation deserves an independent study. Meanwhile, the model focuses on the marker segregation and recombination in a full-sib family, but its principle can also be extended to consider an open-pollinated natural population used to study the genetic structure and evolution of natural populations (Sun et al. 2015).

Our model is based on two-point linkage analysis of fully informative markers. There is still a plenty of room to modify and comprehend the model. First, we assume that an allohexaploid only undergoes a bivalent pairing, but this assumption may be too strong in some situations in which both bivalent and multivalent pairing may occur at the same time. Wu et al. (2004) proposed a mixture model that allows these two types of pairing to be separated in tetraploids from the EM algorithm. More complex EM algorithms should be derived to accommodate to this information in hexaploids. Second, it deserves being extended into three-point linkage analysis because this can not only estimate the combination fraction between two loci, but also examine the influence of genetic interference on the linkage estimation (Wu et al., 2002; Lu et al., 2004; Hou et al., 2009; Liu et al., 2012). Third, subsequent work is needed for QTL mapping to understand the genetic architecture of quantitatively inherited traits in allohexaploids. Despite these extensions being made, our current model provides a general platform to study the linkage and homology of allohexaploids. We have packed the model into computer software at http://ccb.bifu.edu.cn/program.html (available upon the acceptance of this manuscript) which can be freely used by other researchers.

## Acknowledgements

We thank Xiaoming Pang for his contribution to this work and Shandong Research Institute of Pomology for supplying their parsimony data to validate our model. This work is supported by Special Fund for Forest Scientific Research in the Public Welfare (201404102), Changjiang Scholars Award and “Thousand-person Plan” Award.

Authors’ contributions: H.L. wrote the manuscript. X.Z. and Q.Y. derived the model and performed data analysis. Q.Y. and K.M. conducted marker experiment. R.W. conceived of the idea and wrote the manuscript. All authors read and approved the final manuscript.

